# Route of self-amplifying mRNA vaccination modulates the establishment of pulmonary resident memory CD8 and CD4 T cells

**DOI:** 10.1101/2022.06.02.494574

**Authors:** Marco Künzli, Stephen D. O’Flanagan, Madeleine LaRue, Poulami Talukder, Thamotharampillai Dileepan, Andrew G. Soerens, Clare F. Quarnstrom, Sathi Wijeyesinghe, Yanqi Ye, Justine McPartlan, Jason S. Mitchell, Christian W. Mandl, Richard Vile, Marc K. Jenkins, Rafi Ahmed, Vaiva Vezys, Jasdave Chahal, David Masopust

**Affiliations:** Center for Immunology, Department of Microbiology and Immunology, University of Minnesota, Minneapolis, Minnesota, USA; Emory Vaccine Center and Department of Microbiology and Immunology, Emory University School of Medicine, Atlanta, Georgia, USA; Tiba Biotech LLC, Cambridge, Massachusetts, USA; Department of Molecular Medicine, Mayo Clinic, Rochester, Minnesota, USA

## Abstract

Respiratory tract resident memory T cells (Trm), typically generated by local vaccination or infection, can accelerate control of pulmonary infections that evade neutralizing antibody. It is unknown whether mRNA vaccination establishes respiratory Trm. We generated a self-amplifying mRNA vaccine encoding the influenza A virus nucleoprotein that is encapsulated in modified dendron-based nanoparticles. Here we report how routes of immunization in mice, including contralateral versus ipsilateral intramuscular boosts, or intravenous and intranasal routes, influence influenza-specific cell-mediated and humoral immunity. Parabiotic surgeries revealed that intramuscular immunization was sufficient to establish CD8 Trm in lung and draining lymph node. Contralateral, compared to ipsilateral, intramuscular boosting broadened the distribution of LN Trm and T follicular helper cells, but slightly diminished resulting levels of serum antibody. Intranasal mRNA delivery established modest circulating CD8 and CD4 T cell memory, but augmented distribution to the respiratory mucosa. Of note, combining intramuscular immunizations with an intranasal mRNA boost achieved high levels of both circulating T cell memory and lung Trm. Thus, routes of mRNA vaccination influence humoral and cell-mediated immunity, and intramuscular prime-boosting establishes lung Trm that can be further enhanced by an additional intranasal immunization.

## Introduction

Despite the successful control and in some cases eradication of numerous infectious diseases, classical vaccination approaches have failed to eliminate endemic pathogens such as HIV, Malaria, TB, and Influenza, highlighting a need for innovative vaccine design. To combat the emergent threat of SARS-Cov2, novel mRNA vaccine technologies were rapidly developed, adopted, and shown to prevent severe disease outcomes (*1-4*).

Standard immunization includes two intramuscular (IM) immunizations at 21 (BNT162b2) or 28 day (mRNA-1273) intervals, followed by one or more additional boosts 5 months later. Advantages of mRNA vaccination include the relatively potent immunogenicity for CD8 T cells as well as CD4 T cells and antibody, the absence of vector immunity which permits homologous boosting, and the rapidity of production at scale. mRNA vaccines are being considered for diverse disease conditions, including universal influenza vaccines and cancer, both of which likely depend on regional cell-mediated immunity (*5, 6*). This furthers the impetus to better characterize the differentiation and distribution of memory CD8 and CD4 T cells after mRNA vaccination.

mRNA vaccine-elicited T cells have been proposed to contribute to protection upon infection of SARS-Cov2 variants of concern (*7, 8*). In contrast to neutralizing antibodies, T cells recognize partially conserved epitopes and hence may not be completely evaded by mutations in the spike protein of SARS-Cov2. Similarly, T cells recognize epitopes in the nucleoprotein of influenza viruses that may be conserved despite antigenic drift and antigenic shift, unlike neutralizing antibody targets on Hemagglutinin (HA) and Neuraminidase (NA) (*9, 10*). However, mRNA vaccine-elicited T cells have been best characterized in blood, and remain underexplored in the lung, the site of influenza and SARS-Cov2 infections. Indeed, even in mouse studies, little work has been performed on antigen-specific T cells in tissues, including their migration properties, following mRNA vaccination.

Unlike circulating memory T cell populations that patrol blood and lymph, resident memory T cells (T_rm_) are the predominant surveyors of nonlymphoid tissues and accelerate pathogen control in the event of local infection (*11, 12*). Upon reactivation, CD8+ T_rm_ are poised for rapid cytotoxic and innate-like ‘sensing and alarm’ functions which establish an antiviral tissue microenvironment that is further supported by the IFN-y production of CD4+ T_H_1 T_rm_ (*13-16*). In addition, lung resident CD4+ T follicular helper cells (Tfh) support local antibody production after heterosubtypic reinfection (*15, 17*). Trm have also been identified in lung-draining lymph nodes, and local LN memory T cells may make independent contributions to respiratory immunity (*18, 19*). Because of the known protective role of pulmonary Trm in mice and humans, whether mRNA prime-boost vaccination elicits bona fide Trm in the lung or regional LNs is clinically relevant (*16, 19-35*). Induction of pulmonary Trm might represent a supplementary strategy to bolster the efficacy of antibody-targeted vaccines or to induce protective immunity in immunocompromised individuals that have an impaired ability to generate antibodies (*16, 20, 22, 24, 36*). However, previous reports indicate that local antigen expression within the lung is required to establish pulmonary Trm (*20, 23, 37-41*). Therefore, intramuscular mRNA vaccinations may not establish Trm in the respiratory mucosa.

Here, we characterized how antigen-specific CD8 and CD4 T cell and serum antibody responses related to immunization routes after vaccination with modified dendron-based nanoparticle (MDNP)-encapsulated self-amplifying mRNA encoding an influenza nucleoprotein (NP).

## Results

### Impact of ipsilateral versus contralateral intramuscular mRNA-vaccination on CD8 T cell, CD4 T cell, and humoral immunity

To assess memory T cell populations after mRNA vaccination, we primed C57BL/6J mice with 5μg of mRNA encoding the Cal09 influenza nucleoprotein (NP) in the right hamstring. ≥28 days later, we administered an additional 5μg dose of mRNA. To examine whether the administration site of the booster impacted the regionalization of the memory T cell response, we boosted in either the right (ipsilateral) or the left (contralateral) hamstring (Fig. 1A). We used MHC I (H-2K^b^/NP_366-375_) and MHC II (I-A^b^:NP_261-277_) tetramers to track antigen-specific CD8 and CD4 T cell responses in various tissues ≥28 days after the last immunization (Fig S1 A-B) (*42*). Compared to a single immunization, both ipsilateral and contralateral boosting increased the number of H-2K^b^/NP_366-375_+ CD8 memory T cells in the spleen (Fig. 1B). Contralateral immunization was required for establishing memory CD8 T cells in the contralateral lymph node (Fig. 1C-D). This unexpected result might be explained by the lack of CD62L expression on most memory CD8 T cells (Fig. 1C). We also observed that intramuscular (IM) immunization induced a trackable extravascular (IV-neg) CD69^-^ and CD69^+^ H-2K^b^/NP_366-375_^+^ CD8 T cell population in the lung that was bolstered upon boosting (Fig 1E-G).

**Fig 1.**
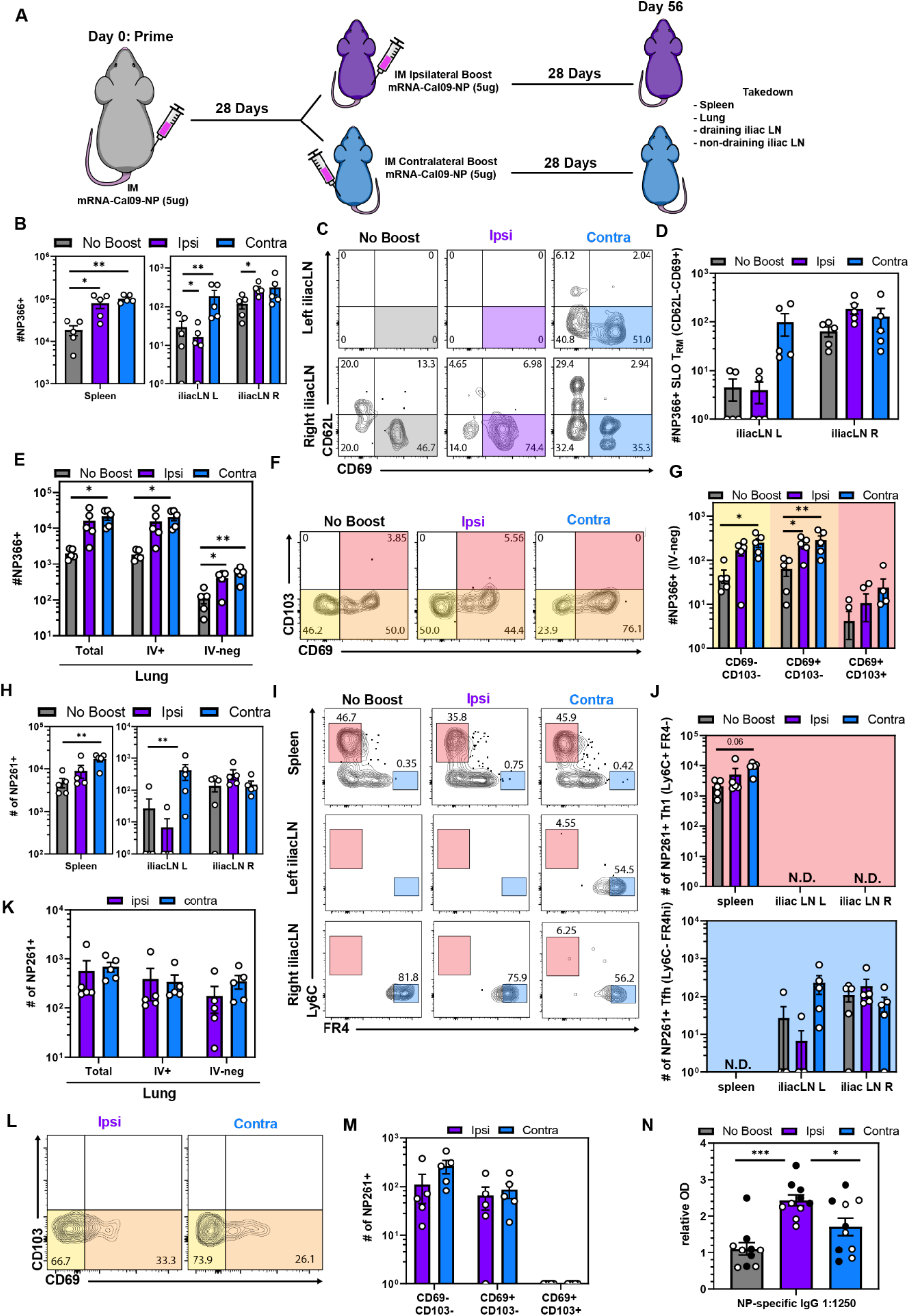
Impact of ipsilateral versus contralateral intramuscular mRNA-vaccination on CD8 T cell, CD4 T cell, and humoral immunity. (A) Mice were primed in the right hamstring then boosted in either the right (ipsilateral) or left (contralateral) hamstring. 28 days post boost, spleen, iliacLN, medLN and lungs were examined. (B) Quantification of antigen-specific CD8+ T cells in the indicated SLOs. (C-D) Representative flow cytometry plot (C) and quantification (D) of NP366-specific CD8+ SLO Trm. (E) Enumeration of antigen-specific CD8+ T cells in the indicated lung compartments. (F-G) Representative flow cytometry plot (F) and quantification (G) of NP366+ CD8 T cell subsets in the lung parenchyma (i.v.-). (H) Enumeration of antigen-specific CD4+ T cells in the indicated SLOs. (I-J) Representative flow cytometry plot (I) and quantification (J) of NP261-specific CD4+ T_H_1 and Tfh in SLOs. (K) Quantification of antigen-specific CD4+T cells in the indicated lung compartments. (L-M) Representative flow cytometry plot (L) and quantification (M) of NP261-specific CD4+ T cell subsets in the lung parenchyma (i.v.-). (N) NP IgG specific serum antibody titers from. Open and closed circles represent data from two independent experiments. Data represent N = 2 independent experiments with n = 4 to 5 mice per group except for (N) were data was pooled from N = 2 independent experiments. Data are shown as mean ± SEM. *P<0.05, **P<0.001, ***P<0.0001 as determined by one-way ANOVA and Tukey’s multiple comparisons test (for 3 groups) or unpaired two-tailed Student’s t test (2 groups). N.D. = not determined.

mRNA prime-boost vaccination also induced regionalization of antigen-specific memory CD4 T cells (Fig 1H). We used the marker combinations Ly6C^+^FR4lo and Ly6C^-^FR4hi to define T_H_1 and Tfh memory subsets, respectively (*43-45*). T_H_1 memory cells were mostly restricted to the spleen, whereas long-lived Tfh were essentially exclusive to the draining lymph nodes (Fig 1 I-J, Fig S1 C-D). Spleen contained a population of cells expressing intermediate levels of FR4 (FR4int). FR4 is also a known T_reg_ marker, however FR4int I-A^b^:NP_261-277_ tetramer+ CD4 T splenocytes do not express Foxp3 (*46*) (Fig. S1E). Intramuscular prime-boost vaccination also induced I-A^b^:NP_261-277_ tetramer+ CD4 memory T cells in the lung parenchyma (Fig 1 K-M). We next investigated the humoral response and found that NP-specific IgG titers were higher in ipsilaterally boosted mice compared to both unboosted and contralateral boosted mice (Fig. 1N). These data indicate that the side of boosting impacts the quality of the immune response, where contralateral boosting established CD69+ CD8 and CD4 Tfh memory T cells in the left iliac LN, but ipsilateral boosting generated slightly higher antibody levels.

### Comparing routes of immunization reveals that intranasal prime-boost vaccination induces more CD103+ CD8 T cells in the lung parenchyma

Human mRNA vaccination is administered by IM injection, though immunization via alternative routes may be an approach to induce specific immunological outcomes (*47*). Conventional RNA lipid nanoparticles (LNPs) cause material-induced inflammatory responses which can cause morbidity and mortality upon intranasal (IN) instillation in mice due to the sensitivity of the respiratory airway to acute inflammation (Fig. S2A-B, Table S1) (*48*). The MDNP formulation selected for this study resulted in complete survival following IN RNA vaccination. Moreover, compared to equivalent RNA doses of conventional LNP, this formula induced negligible material-induced inflammation measured by the induction of IP-10, IL-6, MCP-1, CXCL1, and RANTES after IM immunization (Fig. S2A-C). Given this opportunity, we tested how IM, IN, and intravenous (IV) routes of immunization would compare with respect to memory T cell differentiation and distribution to the respiratory mucosa and lung-draining mediastinal LN (medLN) (Fig. 2A). We observed that the magnitude of H-2K^b^/NP_366-375_ tetramer+ T cells following IM and IV vaccination were largely comparable in secondary lymphoid organs (Fig. 2B). In contrast to IM immunization, IV immunization did not robustly establish CD62L^-^CD69^+^ memory CD8 T cells in the muscle-draining iliac LN (Fig 2C-D). IN vaccination resulted in fewer memory T cells in the spleen and iliac LN but generated more in the medLN (Fig. 2B). After IN vaccination, medLN H-2K^b^/NP_366-375_ tetramer+ T cells phenocopied T cells in the iliac LN of IM immunized mice (Fig. 2C-D). These data demonstrate that vaccination route influences the regionalization and phenotype of mRNA vaccine-elicited T cells.

**Fig 2.**
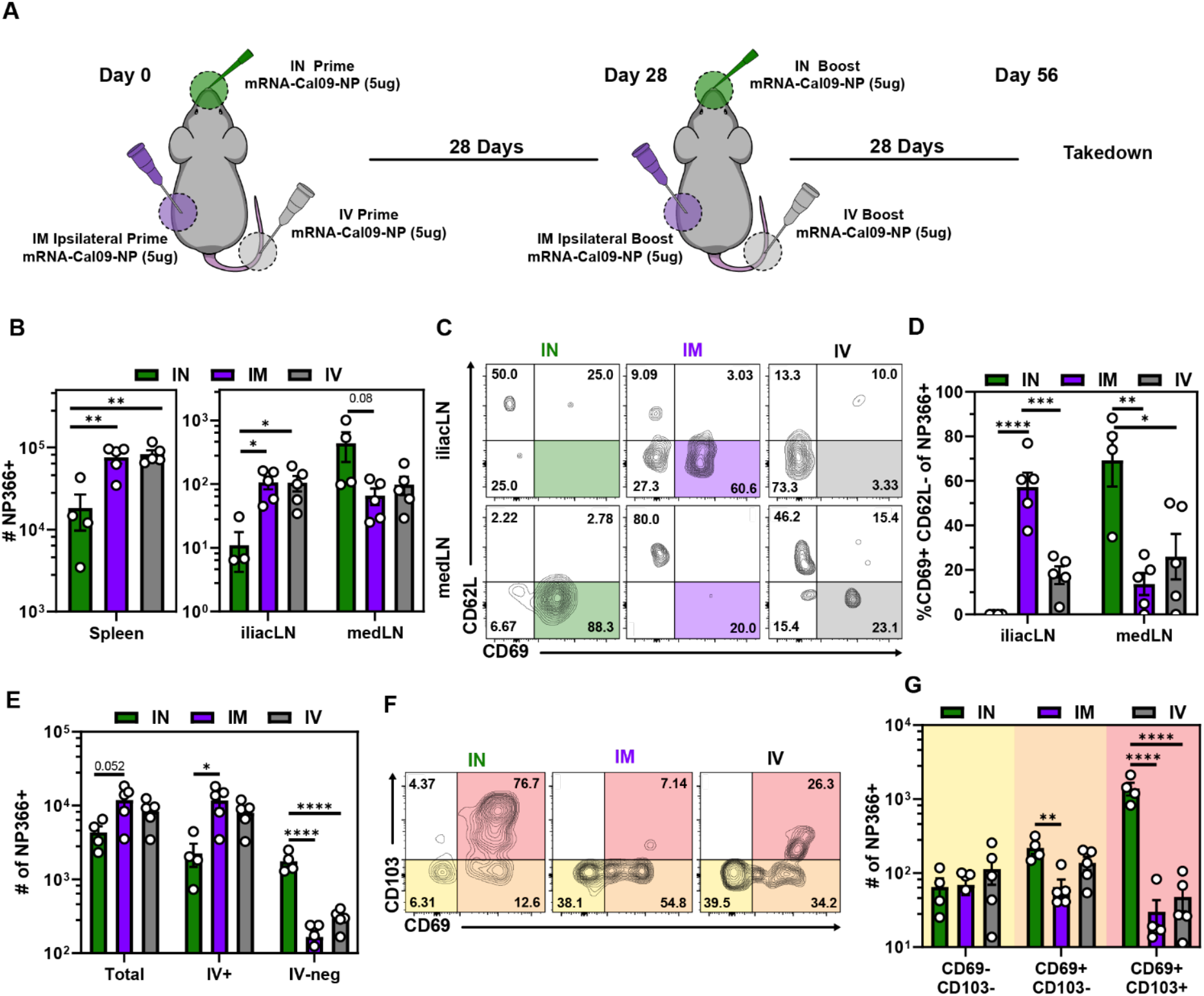
Comparing routes of immunization reveals that intranasal prime-boost vaccination induces more CD103+ CD8 T cells in the lung parenchyma. (A) Mice were either IM, IV or IN prime-boosted and spleen, iliacLN, medLN and lungs were examined 29 days post boost. (B) Quantification of antigen-specific CD8+ T cells in the indicated SLOs. (C-D) Representative flow cytometry plot (C) and quantification (D) of NP366-specific CD8+ SLO Trm. (E) Quantification of antigen-specific CD8+ T cells in the indicated lung compartments. (F-G) Representative flow cytometry plot (F) and quantification (G) of NP366+ CD8 T cell subsets in the lung parenchyma (i.v.-). Data represent N= 2 independent experiments with n = 4 to 5 mice per group. Data are shown as mean ± SEM. *P<0.05, **P<0.001, ***P<0.0001 as determined by one-way ANOVA and Tukey’s multiple comparisons test.

While IM and IV vaccinations induced demonstrable T cells within the extravascular compartments of the lung, most cells were labeled with iv antibody. In comparison, IN prime-boost vaccination established ∼10x more extravascular NP366+ T cells within the lung (Fig. 2E). Unlike IM and IV elicited CD8 T cells, most influenza-specific memory CD8 T cells following IN mRNA administration were CD69^+^CD103^+^, a phenotype that was previously associated with localization in proximity to airways (Fig. 2F-G) (*18, 20, 49-51*). Together, these data highlight that all routes of immunization established broadly distributed memory T cells, but that intranasal immunization increased the magnitude of CD69^+^CD103^+^ CD8 T cells in the lung and medLN.

### mRNA vaccination generates long-lived Tfh in the draining LN but not in the lung

The impact of intranasal mRNA delivery on the CD8 memory compartment prompted us to investigate whether the route also impacted antigen-specific CD4 memory T cell magnitude and phenotype in the lung and draining lymph node (Fig. 3A). In contrast to CD8 T cells, the magnitude of antigen-specific CD4 T cells in the spleen following IN or IM vaccination was comparable, whereas the abundance of I-A^b^:NP_261-277_+ T cells in the different lymph nodes correlated with the vaccination route (Fig. 3B). Again, T_H_1 cells were enriched in spleen and Tfh were restricted to the draining lymph node (Fig. 3 C-D, Fig. S3A-B). NP-specific IgG serum antibody titers were not affected by vaccination route (Fig. 3E). We observed a significant increase in I-A^b^:NP_261-277_+ CD4s in the lung after IN vaccination and this was completely accounted for by an increase in extravascular cells (Fig. 3F), many of which were CD69^+^ (Fig. 3G-H). In contrast to NP366+ CD8 memory T cells, CD103 upregulation was moderate, consistent with previously reported phenotypes of CD4 T cells in non-lymphoid tissues (*52*). In addition to well-described pulmonary T_H_1 resident CD4 T cells (PSGL1hi FR4lo), a Tfh-like memory population (PSGL1lo FR4hi) was recently identified in the lung following influenza infection that promoted local antibody production upon reactivation (*14, 15*). In contrast to influenza infection, mRNA vaccination did not induce I-A^b^:NP_261-277_+ PSGL1lo FR4hi cells in the lung (Fig. 3I-J, Fig. S3C). However, IN vaccination reproducibly induced an FR4 intermediate population that otherwise phenocopied T_H_1-like resident memory T cells (Fig. S3D) (*14, 15*). Taken together, IN vaccination induced long-lived Tfh in the medLN and in contrast to IM vaccination generated a robust pulmonary CD4 T cell memory population.

**Fig 3.**
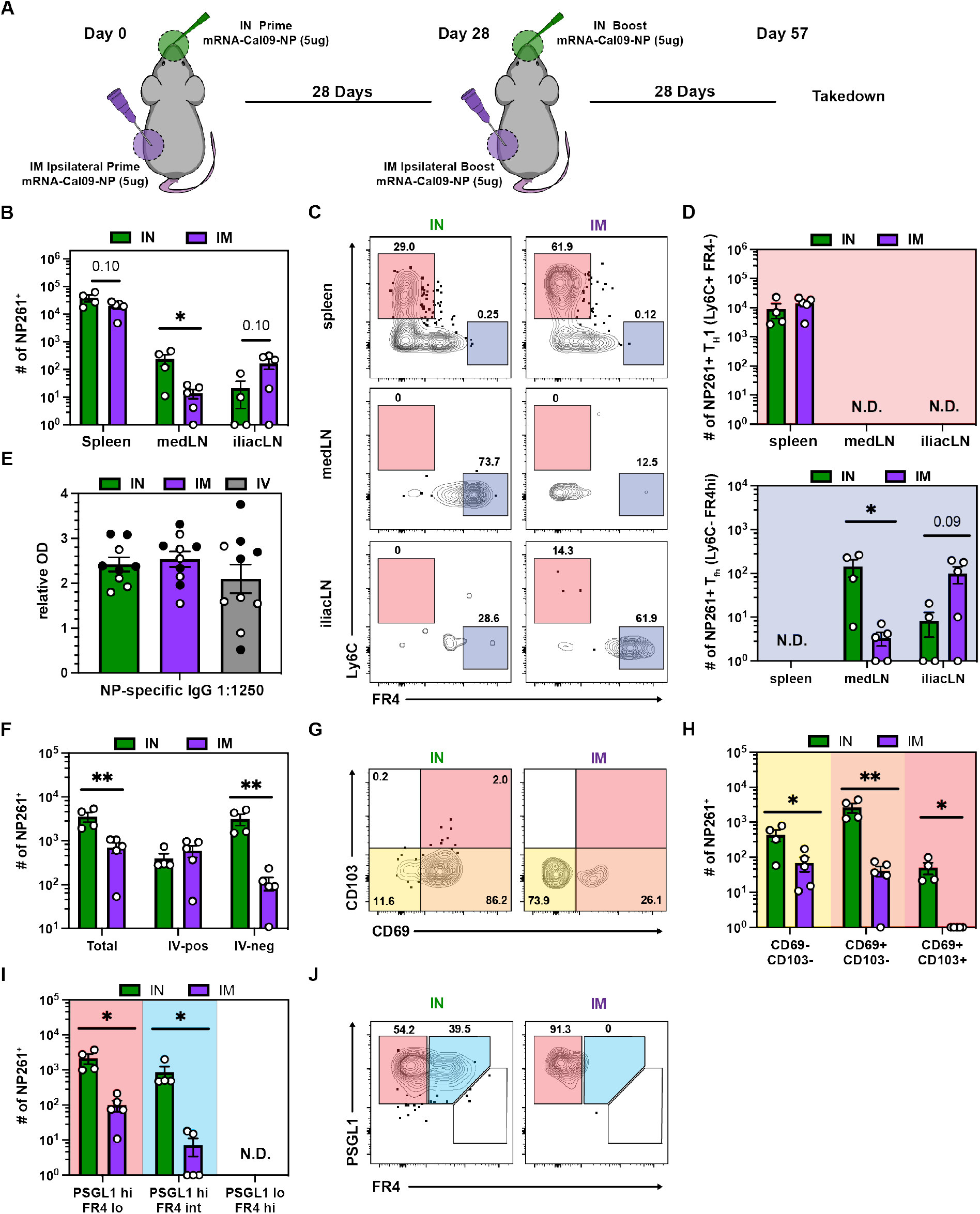
mRNA vaccination generates long-lived Tfh in the draining LN but not in the lung. (A) Mice were either IM or IN prime-boosted and spleen, iliacLN, medLN and lungs were examined 29 days post boost. (B) Quantification of antigen-specific CD4+ T cells in the indicated SLOs. (C-D) Representative flow cytometry plot (C) and quantification (D) of NP261-specific CD4+ T_H_1 and Tfh in SLOs. (E) NP IgG specific serum antibody titers. Open and closed circles represent data from two independent experiments. (F) Quantification of antigen-specific CD4+ T cells in the indicated lung compartments. (G-H) Representative flow cytometry plot (G) and quantification (H) of NP261-specific CD4+ T cell subsets in the lung parenchyma (i.v.-). (I-J) Quantification (I) and representative flow cytometry plot (J) of NP261-specific CD4+ T cell subsets in the lung parenchyma (i.v.-). Data represent N= 2 independent experiments with n = 4 to 5 mice per group. Data are shown as mean ± SEM. *P<0.05, **P<0.001, ***P<0.0001 as determined by one-way ANOVA and Tukey’s multiple comparisons test (for 3 groups) or unpaired two-tailed Student’s t test (2 groups). N.D. = not determined.

### All mRNA immunization routes were sufficient to induce pulmonary resident memory CD8 T cells

Upon IM mRNA prime-boost vaccination, we noted a population of CD8 and CD4 memory T cells in the lung, some of which expressed CD69. To further validate the presence of H-2K^b^/NP_366-375_ tetramer+ CD8 T cells in the lung after intramuscular immunization, we performed *in situ* tetramer staining and found influenza-specific memory T cells in the lung parenchyma (Fig. 4A). Although CD69 is often used as a proxy to denote resident memory T cells, many Trm do not express CD69 and not all CD69^+^ cells are resident (*11, 53*). Moreover, the establishment of Trm in the lung often requires local antigen recognition (*20, 23, 37-41*). Therefore, we wondered whether or not these cells were resident or recirculating. To assess migration of mRNA vaccine-elicited pulmonary T cells, we conjoined CD45.2+ mRNA primed-boosted mice to congenically mismatched naive CD45.1+ C57BL/6J mice via parabiosis surgery (Fig. 4B). While circulating bloodborne populations equilibrate between the two mice, the failure of cells to equilibrate in corresponding tissues is strong evidence for residency. Three weeks after surgery, the blood contained equal proportions of CD45.2+ and CD45.1+ CD8 T cells, demonstrating successful anastomosis between the immune and naïve parabionts (Fig. 4C). To calculate the proportion of resident T cells, we compared the number of antigen-specific T cells in the immune versus naive parabiont using a previously reported equation (*11*). Unexpectedly, extravascular pulmonary NP366+ CD8 T cells were preferentially present in the immune parabiont – demonstrating that IM mRNA vaccination elicits bona fide resident memory T cells (Fig. 4D-E). The few NP366+ CD8 T cells found in the lung of the naive parabiont were >90% CD69^-^ CD103^-^. The proportion of resident cells increased stepwise from 40% in the CD69^-^CD103^-^ compartment to 95% and 100% residence in the CD69^+^CD103^-^ and CD69^+^CD103^+^ subsets respectively. To test whether local antigen deposition impacts the formation of pulmonary Trm, we repeated parabiosis with IN prime-boosted mice. Not only did IN vaccination boost the number of extravascular pulmonary T cells by ∼10 fold, the fraction of resident T cells also increases from ∼70% to ∼90% (Fig. 2E, Fig. 4 D-E). We also assessed residence within the CD4 compartment. Our analysis was limited to the IN vaccination route due to insufficient recovery of antigen-specific CD4 T cells upon parabiosis after IM vaccination. Following IN mRNA vaccination, ∼90% of I-A^b^:NP_261-277_+ CD4 T cells resided in the lung for at least three weeks and this proportion increased to 100% if the analysis was limited to CD69^+^ cells (Fig. 4 F-G). In conclusion, mRNA vaccination induces bona fide Trm independent of vaccination route.

**Fig 4.**
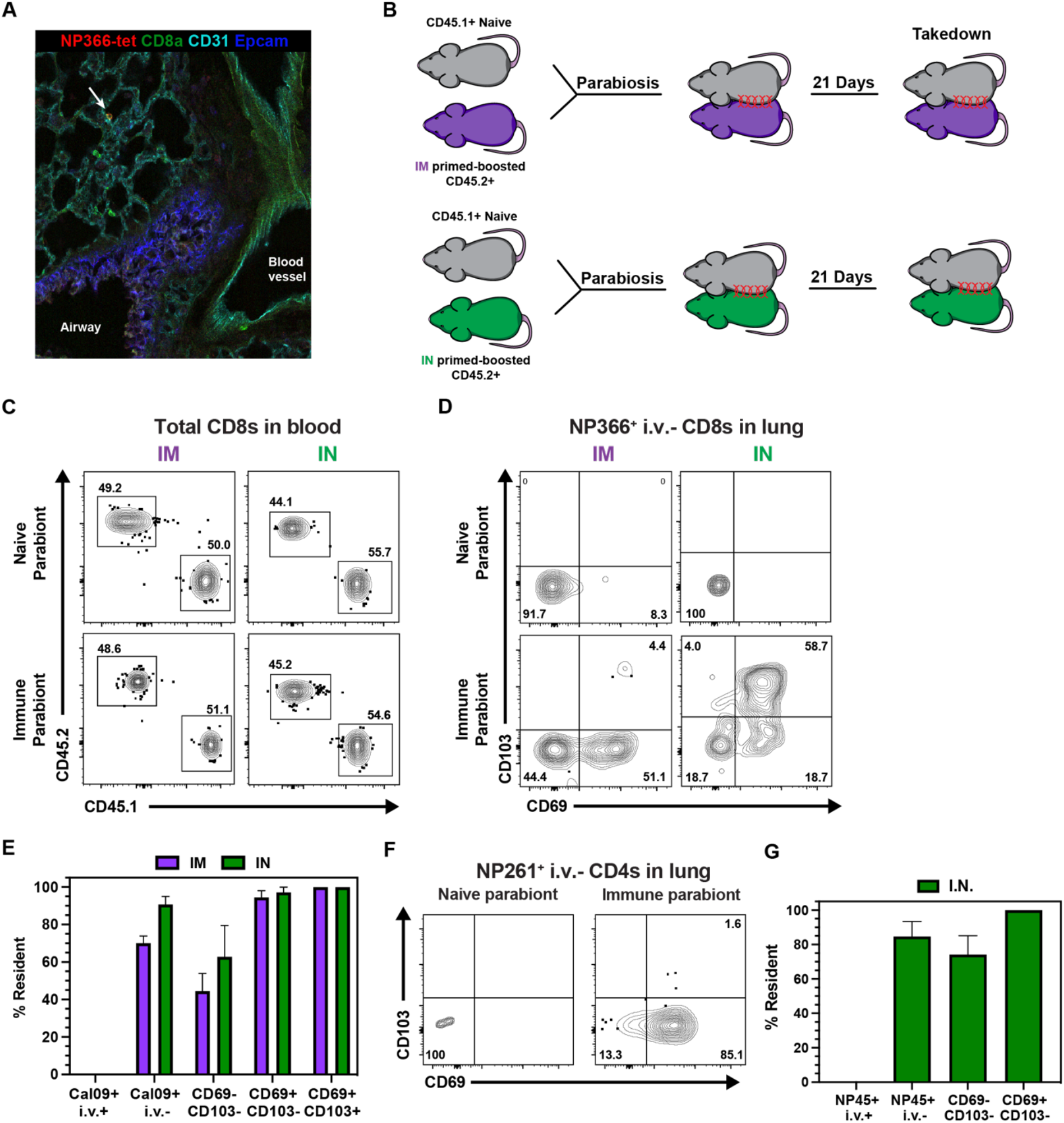
All mRNA immunization routes were sufficient to induce pulmonary resident memory CD8 T cells. (A) Representative image of in situ NP366-tetramer staining in the lung upon intramuscular prime-boost. (B) CD45.2+ mice were either IM or IN prime-boosted and then conjoined to CD45.1+ naïve hosts. After 21 days, lungs were harvested to examine recirculating and resident populations antigen-specific CD8 and CD4 T cells. (C) Representative flow cytometry plots depicting CD45.1+ and CD45.2+ CD8 T cells to demonstrate equilibration in the blood. (D-E) Representative flow cytometry plot (D) and quantification (E) of NP366-specific CD8 T cell subsets in the lung parenchyma. (F-G) Representative flow cytometry plot (F) and quantification (G) of NP261-specific CD4 T cell subsets in the lung parenchyma. Data represent N= 2 independent experiments with n = 4 to 5 mice per group, except for (A) which represents N=1 independent experiment with n = 3 mice. Data are shown as mean ± SEM. *P<0.05, **P<0.001, ****P<0.0001 as determined by unpaired two-tailed Student’s t test with N= 2 independent experiments.

### Combining intramuscular immunizations with an intranasal mRNA boost achieves high levels of both circulating memory and lung Trm

While IM vaccination resulted in a large population of NP366+ CD8 T cells in the circulation, IN immunization biased differentiation towards a higher magnitude of pulmonary Trm. We next tested if 1) antigen-specific memory T cells retain the potential to generate abundant lung Trm populations and 2) whether one could generate a large pool of both circulating and pulmonary T cells by giving IM prime-boosted mice a secondary IN boost (Fig 5A). Compared to IM prime-boost alone, IN boosting increased the number of antigen-specific CD8 and CD4 T cells in the spleen, medLN, and lung (i.v.-), but not in the iliac LN (Fig. 5B-C). IN boosting expanded the proportion and number of CD69^+^CD103^+^ CD8 T cells and CD69^+^ CD4 T cells in the extravascular compartment of the lung, comparable to the IN prime-boost regimen (Fig 2E, 3F, Fig.5. D-G). These data demonstrate that combining intramuscular prime-boost immunizations with a third intranasal mRNA immunization achieves high levels of both circulating and lung memory T cells.

**Fig 5.**
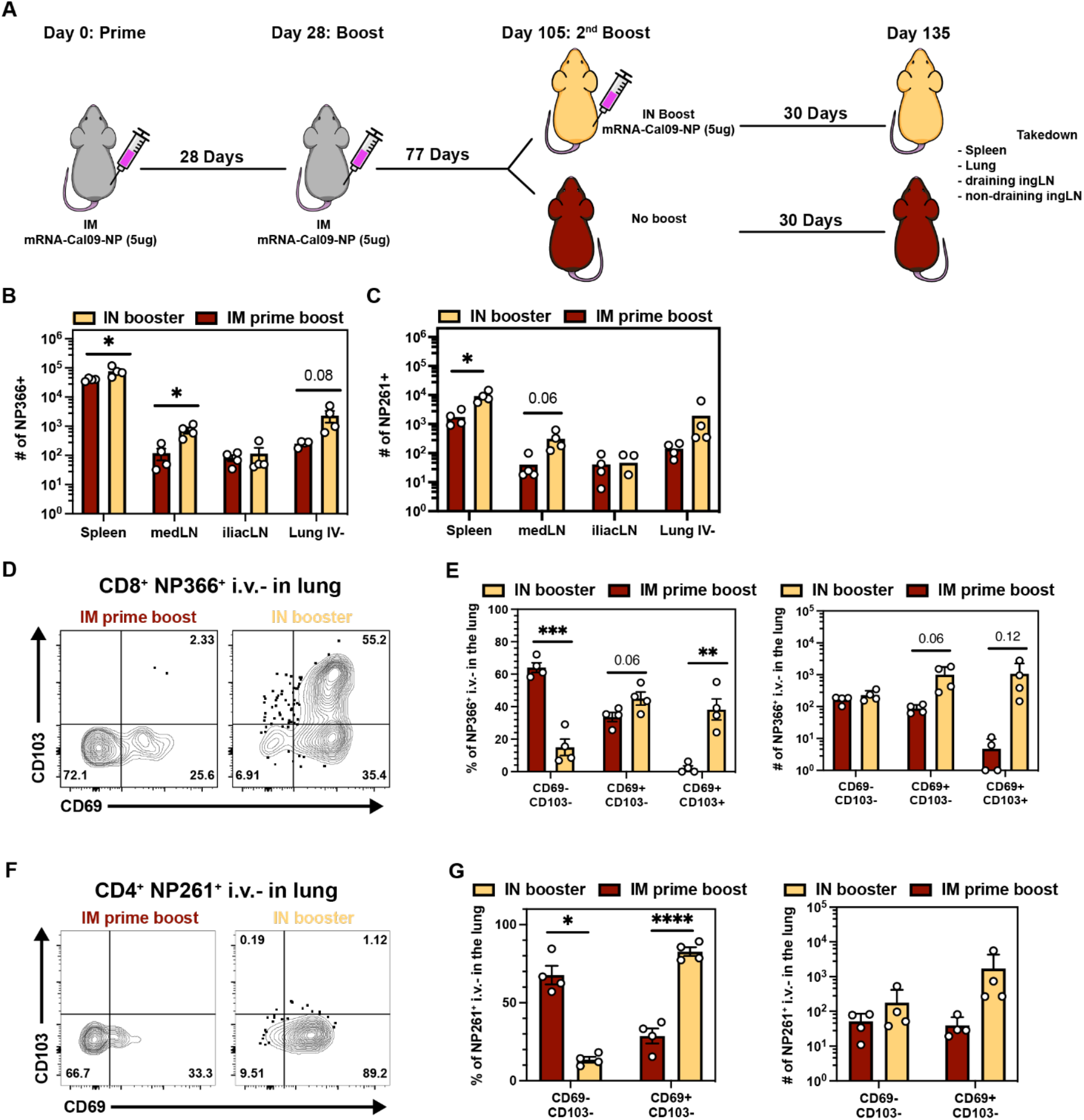
Combining intramuscular immunizations with an intranasal mRNA boost achieves high levels of both circulating memory and lung Trm. (A) IM prime-boosted mice received a second booster via intranasal route and spleen, iliacLN, medLN and lungs were examined 28 days post secondary boost. (B-C) Quantification of antigen-specific CD8+ T cells (B) and CD4+ T cells (C) in the indicated compartments. (D-E) Representative flow cytometry plot (D) and quantification (E) of i.v.- NP366-specific CD8+ T cell subsets in the lung. (F-G) Representative flow cytometry plot (F) and quantification (G) of i.v.- NP261-specific CD4+ T cell subsets in the lung. Data represent N= 2 independent experiments with n = 4 to 5 mice per group. Data are shown as mean ± SEM. *P<0.05, **P<0.001, ****P<0.0001 as determined by unpaired two-tailed Student’s t test with N = 2 independent experiments.

## Discussion

In this study, we characterized the abundance, phenotype, and anatomic distribution of mRNA vaccine-elicited antigen-specific CD8 and CD4 memory T cells (Fig. S4). Independent of route, mRNA vaccination induces pulmonary Trm, where frontline cellular immunity is known to contribute to protection against certain respiratory infections (*16, 19-35*). Thus, mRNA vaccines may fulfill an important criterion in the pursuit of a universal influenza vaccine: establishing cellular immune memory at the primary site of infection. Importantly, we found that prime-boosted memory T cells elicited by intramuscular vaccinations retain the potential to generate abundant pulmonary Trm upon an additional IN boost. These data could inform a strategy to leverage pre-existing memory T cells to bolster frontline immunity at the site of infection without compromising the circulating compartment. This could have implications for future iterations of SARS-CoV2 vaccination, or other respiratory pathogens. Including conserved T cell epitopes in vaccines that also establish humoral immunity may provide immunity in immunocompromised individuals that have impaired ability to efficiently generate antibodies or against pathogens that evolve to evade vaccine-elicited neutralizing antibodies. With that said, this study focused primarily on the differentiation and distribution of mRNA vaccine-elicited memory CD4 and CD8 T cells. Future studies will be needed to integrate the quality of cell-mediated immunity with protective potential. This is a complex issue, and will depend on the route, dose, virulence, and antigenic profile of the pathogens in question, as well as availability of animal models that adequately predict human outcomes.

Lazcko and colleagues demonstrated the presence of activated CD69^+^ IV-neg T cells in the lung 10 days after IM mRNA-LNP vaccination and a recent preprint from Mao et al. shows persistence of antigen-specific CD8 T cells in the lung for at least 56 days, raising the question of whether Trm are established (*54, 55*). Here, we report the persistence of vaccine-elicited antigen-specific CD8 and CD4 T cells in the lung for at least 135 days after IM priming. Numerous studies have shown that establishment of pulmonary CD8 Trm requires local antigen recognition (*20, 23, 37-41*). To our surprise, based on migration assays in parabiotic mice most CD8 memory T cells, including CD69^-^ cells, were Trm. Although we would not expect antigen to be present in the lung after IM immunization, we cannot exclude this possibility. Nevertheless, intranasal immunization was much more effective at establishing pulmonary Trm compared to IV and IM immunization.

We found it surprising that IN and IV immunizations were effective at establishing memory CD8 and CD4 T cells, as the vaccine is tailored for intramuscular administration. A previous report of self-amplifying RNA delivery by the IN route using multiple delivery systems including LNPs showed poor immunogenicity relative to more typical IM or intradermal injections, and was associated with rapid clearance (*56*). For the most part, intranasal delivery of mRNA vaccines have been attempted using well-established polymer-based transfection reagents, though such polymer-based approaches have not historically reached clinical application successfully, possibly in part due to toxicity issues (*57-60*). Moreover, IN LNP can induce lethal inflammatory responses in mice, complicating studies of alternative administration routes (*48*). The MDNP formulation here did not have that effect and induced reduced level of proinflammatory cytokines. It is unclear whether this could be translated to humans. Of note however, we found that IN immunization of mice that had previously received two IM immunizations resulted in particularly robust memory CD8 T cells in both the pulmonary and circulating compartments. CD4 T cells were augmented in lung as well. These data add to the possibility that future mRNA vaccinations, perhaps combined with systemic immunizations, might be tailored to focus immunity at specific sites including the respiratory mucosa based on modulating the site of boosting.

A common question concerning mRNA vaccines is whether one should get a booster vaccine in the same (ipsilateral) or opposite (contralateral) arm. In the mouse model, we found moderate differences in the immunological parameters that we measured. Serum antibody titers were higher after ipsilateral versus contralateral boosting. A recent report indicated that affinity maturation was also enhanced by same-site boosting, suggesting that ipsilateral boosting may offer advantages for humoral immunity (*61*). Opposite-site boosting readily affected T cell responses. Tfh were limited to LNs that drained the site of immunization (and absent from both spleen and lung), so exhibited broader distribution after contralateral immunization. This observation also extended to CD69^+^CD62L^-^ memory CD8 T cells, a phenotype previously shown to correlate with LN residents (*53*). LN Trm putatively derive from retrograde migration from upstream nonlymphoid tissues, so this population may reflect ex-Trm derived from muscle (*18*). However, preliminary attempts to examine muscle tissue did not yield viable lymphocyte populations that could be analyzed. We also observed that Tfh and Trm-phenotype CD8 T cells were uniquely established in the lung draining LN after IN immunization.

RNA-based vaccines are rapidly scalable against essentially any protein antigen and are not vulnerable to vector immunity that could otherwise limit opportunities for boosting or reusing the same vector, encoding different antigens, in future vaccines. The prospects are exciting for combatting emerging pathogens, antigenically variable pathogens that might be addressed by megavalent vaccination, and perhaps non-infection conditions such as personalized tumor vaccines. For these reasons, it is important to more fully characterize RNA vaccine immunogenicity. The establishment of Trm is of obvious importance, but difficult to assess in humans, despite COVID-19 providing intense interest and subject availability. Genetically defined mice and parabiosis surgery provided an opportunity to address this fundamental question and suggest that RNA vaccines may be effectively exploitable for establishing Trm.

## Material and Methods

### Study design

The aim of this study was to characterize the self-amplifying mRNA-MDNP vaccine-elicited CD8 and CD4 T cell responses against the Cal09 nucleoprotein of the Influenza virus in secondary lymphoid organs and the lung. We used major histocompatibility complex I and II tetramers to identify polyclonal antigen-specific CD8 and CD4 memory T cells. Analysis of experimental data was conducted unblinded. Detailed description of experimental replicates, sample sizes, and statistical analysis can be found in the figure legends or under the section “Statistical analysis”.

### Mice

C57BL/6J (B6) and B6.SJL-Ptprc^a^Pepcb/BoyJ (B6 CD45.1), were purchased from Jackson Laboratory and maintained under specific pathogen-free conditions at the University of Minnesota according to the Institutional Animal Care and Use Committees guidelines. All B6 mice used in the experiments were female and 6-9 weeks of age at the timepoint of the first immunization. For inflammatory response testing, BALB/cJ mice were purchased from Jackson Laboratory and maintained according to Tiba Biotech’s Institutional Animal Care and Use Committee guidelines. All ALB/cJ mice were 6-10 weeks of age at time of experimentation.

### MDNP vaccine production

The Cal/09 nucleoprotein coding sequence was codon-optimized and cloned into a DNA plasmid encoding a self-amplifying mRNA template based on the genome of Venezuelan equine encephalitis virus, followed by RNA synthesis by run-off in vitro transcription from an upstream T7 promoter and subsequent enzymatic capping essentially as reported previously (*62*). Expression potency of each lot of self-amplifying mRNA produced for this work was validated by transfection of BHK cells and subsequent immunoblot using anti-Influenza A Virus Nucleoprotein (NP) antibody (Clone 1C5-1B7 NR-43899) obtained through BEI Resources, NIAID, NIH, (data not shown). The capped self-amplifying RNA was dissolved in 10mM citrate buffer and mixed using a NanoAssemblr Ignite microfluidic mixer (Precision Nanosystems) with a proprietary ethanolic solution of an ionizable modified dendron provided by Tiba Biotech (at an N:P ratio of 4), plus cholesterol, DOPE, and DMG-PEG2000 (Avanti Polar Lipids) in a 100:288:60:1 molar ratio respectively. Nanoparticles were dialyzed against sterile, endotoxin-free PBS using 20,000 Da molecular weight cutoff dialysis cassettes and sterile filtered using 0.2 micron poly(ether sulfone) filters (CELLTREAT Scientific Products). Nanoparticle diameter and polydispersity index were assessed by dynamic light scattering and encapsulation efficiency measured by RiboGreen® (Thermo) dye-exclusion assay. Representative nanoparticle characteristics of the final Cal/09 nucleoprotein self-amplifying mRNA vaccine are provided in Table S2.

### Immunizations

Mice were immunized with 5μg (50μl at 0.1mg/ml) of mRNA encapsulated in MDNP encoding for the nucleoprotein of the Influenza A H1N1 Cal/09 strain per dose. Vaccine was either delivered intramuscular (right or left hamstring), intranasal, or intravenous. For injections, mice were either anaesthetized with vaporized isoflurane (intramuscular) or ketamine and xylazine (intranasal).

### Lymphocyte isolation from spleen, lymph nodes, and lung

Mice were injected intravenously (tail vein) 15 minutes prior to sacrifice with NAD-induced cell death (NICD) protector (25 μg, Biolegend, #149802) to prevent NAD-induced cell death during tissue processing (*45*). Intravascular staining was used to discriminate cells present in the vasculature from cells present in the tissues parenchyma as previously described (*63*). Briefly, 3 minutes prior to euthanasia mice were injected retroorbitally with 3µg of fluorescently conjugated αThy1.2 antibody (BV650, BD, #740443). Lymphocyte isolation from spleen, LNs, and lung was performed as described (*11*). Spleen and LN single cell suspensions were generated via dissociation through 70µm filters (Falcon, #352350). For lung cell isolations, lungs were excised, minced, and digested in Collagenase I (Worthington, #LS004197) for 1 hour at 37°C shaking at 200rpm. Lung homogenates were then mechanically dissociated using a gentleMACS Dissociator (Miltenyi Biotec) and subsequently passed through a 70µm filter. Finally, lymphocytes were enriched on a Percoll gradient. Cell enumerations were determined using PKH26 Reference Microbeads (Sigma-Aldrich, # P7458-100ML) as previously described (*64*).

### Flow cytometry

To identify influenza-specific CD8+ T cells, isolated lymphocytes were surface-stained with homemade PE-conjugated H-2K^b^/NP_366-375_ tetramer for 1 hour at room temperature in the presence of viability dye (Ghost Dye Red 780, Tonbo Biosciences, #13-0865-T500), Dasatinib (50nM), Fc Shield (Tonbo, #70-0161-M001), and indicated antibodies. Splenic influenza-specific CD4+ T cells were isolated by staining single cell suspensions with homemade APC-conjugated I-A^b^:NP_261-277_ tetramer for 1 hour at room temperature in the presence of Dasatinib (50nM) followed by tetramer enrichment (*64, 65*). CXCR5 (BV421, BioLegend, #145512) was added at the time of tetramer staining. Further surface staining was performed for 30 minutes on ice in the presence of viability dye (Ghost Dye Red 780, Tonbo Biosciences, #13-0865-T500). Lung and LN-derived I-A^b^:NP_261-277_ tetramer+ CD4 T cells were identified by staining single cell suspensions with I-A^b^:NP_261-277_ APC tetramer for 1 hour at room temperature in the presence of viability dye (Ghost Dye Red 780, Tonbo Biosciences, #13-0865-T500), Dasatinib (50nM), Fc Shield (Tonbo, #70-0161-M001), and indicated antibodies. All stained samples were acquired with LSR Fortessa flow cytometers (BD) and analyzed with FlowJo software (TreeStar).

Isolated H-2K^b^/NP_366-375_ tetramer+ lymphocytes were stained with the following antibodies: CD8a (BUV496, BD, #750024), CD45.2 (AF700, BioLegend, #109822), CD69 (BV421, BD, #562920), CD103 (BV510, BD, #563087), CD44 (BV785, BioLegend, #103041), CD62L (BUV737, BD, #612833), and were indicated CD45.1 (APC, BioLegend, # 110714)

Isolated I-A^b^:NP_261-277_ tetramer+ lymphocytes were stained with the following antibodies: CD4 (BUV496, BD, #612952), CD45.2 (AF700, BioLegend, #109822), CD103 (BV510, BD, #563087), CD44 (BV785, BioLegend, #103041), CD62L (BUV737, BD, #612833), Ly6C (FITC, BD, #553104), CD69 (PE-Dazzle, BioLegend, #104536), FR4 (PE-Cy7, eBioscience, #25-5445-82), PSGL1 (BV605, BD, #740384), CXCR6 (BV711, BioLegend, # 151111), PD-1 (BUV395, BD, #744549), and were indicated CD45.1 (PE, BD, # 553776).

### Enzyme-linked immunosorbent assay (ELISA)

96 well plates (ThermoFisher, #44-2404-21) were coated with the Cal09 influenza nucleoprotein (1μg/m, Sino Biological, #40205-V08B-100) and incubated overnight at 37°C. Using the Invitrogen ELISA Kit (#88-50400-88), plates were blocked for 2 hours at 37°C before diluted sera were added and incubated for 1 hour at 37°C. HRP-conjugated goat anti-mouse IgG detection antibody (Jackson Immuno, #115-035-062) was incubated for 1 hour at 37°C before plates were developed with 1x TMB for 15 min followed by addition of stop solution (Biofix, #NC9141468). Optical density was measured at 450nM wavelength using a Tecan Infinite M Plex.

### Immunohistochemistry

Lung tissues were manually cut into sections using surgical blades and incubated with PE-conjugated-H-2K^b^/NP_366-375_ overnight at 4°C. Sections were fixed in 2% paraformaldehyde in PBS for 2 hours at 4°C and incubated in 30% sucrose overnight at 4°C before freezing in Tissue-Tek O.C.T. compound. 20μM thick sections were generated using the Leica Cryostat and subsequently stained with CD8 BV480 (BD, #566096), CD31 AF488 (Biolegend, #102514), Epcam AF594 (Biolegend, #118222) and anti-PE (Novusbio, #NB120-7011). Tetramer staining was further amplified using Cy3-donkey anti-rabbit (Jackson Immuno, #711-165-152).

### Parabiosis surgery

Parabiosis surgeries were performed as previously described (*13*). In brief, mice were anaesthetized with Ketamine and Xylazine and fur was depilated on each mouse along the opposite lateral flank via electric clippers. The skin was wiped clean and sterilized with alcohol prep pads, Betadine solution, and 70% alcohol. Identical incisions were made on the lateral aspect of each mouse from the hip to shoulder and 9mm wound clips (BD, #427631) were used to approximate and join the dorsal and ventral skins of adjacent mice. Conjoined mice were then allowed to rest for 21 days before sacrifice and analysis. Equilibration was confirmed in the peripheral blood prior to euthanasia.

### Statistical analysis

Data were analyzed and visualized using Prism 9 (GraphPad). Statistical significance was determined by one-way ANOVA and Tukey’s multiple comparisons test (more than two groups) or unpaired two-tailed Student’s t test (two groups) with P < 0.05, **P < 0.01, ***P < 0.001, ****P < 0.0001. Data are shown as mean ± SEM as indicated in figure legends.

## Acknowledgments

We thank the Masopust and Vezys group members for helpful discussions. Funding: 75N93019C00051 Collaborative Influenza Vaccine Innovation Centers (CIVICs) (D.M., R.A.), Minnesota Partnership for Biotechnology and Medical Genomics (D.M., M.B., R.V.), National Institutes of Health grant 3R01AI084913 (D.M.), Swiss National Science Foundation (SNSF) grant P2BSP3_200187 (M.K.)

## Author contributions

M.K., S.O., M.L. M.B., R.V., Y.Y., R.A., V.V. and D.M. conceptualized the project. M.K., S.O., M.L., and D.M. designed the experiments. M.K., S.O., A.G.S, C.Q., S.W., and J.S.M. performed experiments. M.K., S.O. analyzed and visualized the data. M.K., S.O., and D.M. wrote the manuscript. P.T., J.M., C.W.M., J.C., T.D., and M.J. designed and provided resources.

## Competing interests

P.T, J.M., C.W.M., J.C. work for Tiba Biotech LLC which is a for-profit company.

## Supplementary Material

**Table S1.** Clinical scoring rubric for inflammation and respiratory distress. *Humane endpoints: If a mouse has either an Appearance or Respiration score of 4, the mouse will be euthanized. If a cumulative score of >4 is observed, the mouse will be euthanized*.

| Variable | Score and description |
| --- | --- |
| **Appearance/Attitude** | 0- active, smooth coat |
|  | 1- mild hunched posture +/- piloerection |
|  | 2- moderate hunched posture, “puffy” from piloerection |
|  | 3- decreased body condition (thin) +/- hunched posture, piloerection, dehydration based on low skin elasticity from dorsal pinch test |
|  | 4- Low activity level (decreased responsiveness to stimuli), decreased body condition +/- piloerection |
| **Respiration quality** | 0- Normal |
|  | 1- Increased respiratory rate, normal effort |
|  | 2- Increased respiratory rate, some effort (abdominal) |
|  | 3- Increased abdominal effort, intermittent gasping |
|  | 4- High effort, Gasping |

**Table S2.**
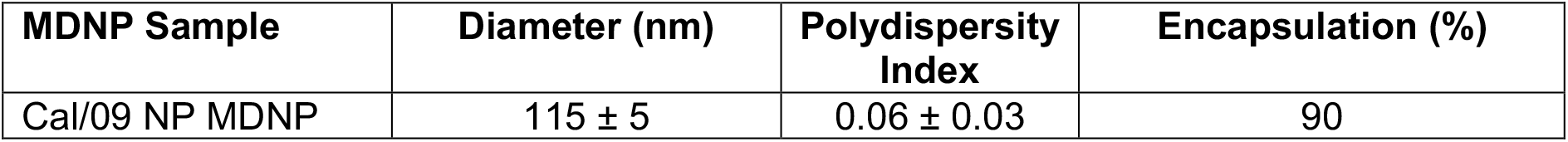
Nanoparticle size, polydispersity, and encapsulation efficiency. Average of 3 independent measurements ±SD shown for diameter and PDI as measured by DLS.

**Fig. S1:**
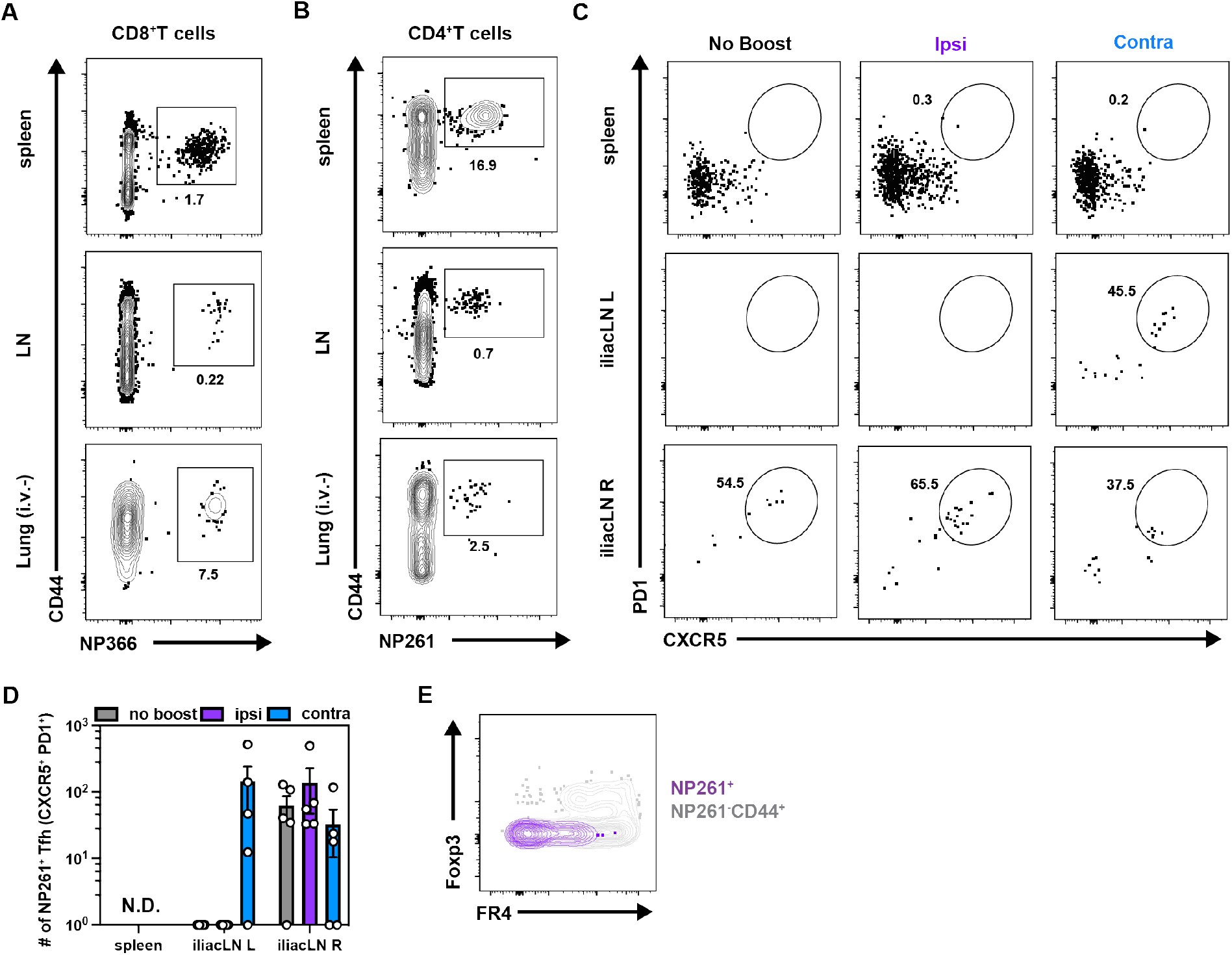
Impact of ipsilateral versus contralateral intramuscular mRNA-vaccination on CD8 T cell, CD4 T cell, and humoral immunity. (A) Representative flow cytometry plot of CD8 T cell NP366^+^ tetramer staining in various tissues. (B) Representative flow cytometry plot of CD4 T cell NP261^+^ tetramer staining in various tissues. (C-D) Representative flow cytometry plot (C) and quantification (D) of NP261-specific CD4+ Tfh identified by high expression of PD1 and CXCR5 in SLOs. (E) Representative flow cytometry plot of NP261+ CD4+ T cells (grape) and endogenous IV-CD44+ CD4 T cell compartment (grey). Data represent N= 2 independent experiments with n = 4 to 5 mice per group. Data are shown as mean ± SEM. *P<0.05, **P<0.001, ***P<0.0001 as determined by unpaired two-tailed Student’s t test. N.D. = not determined.

**Fig S2.**
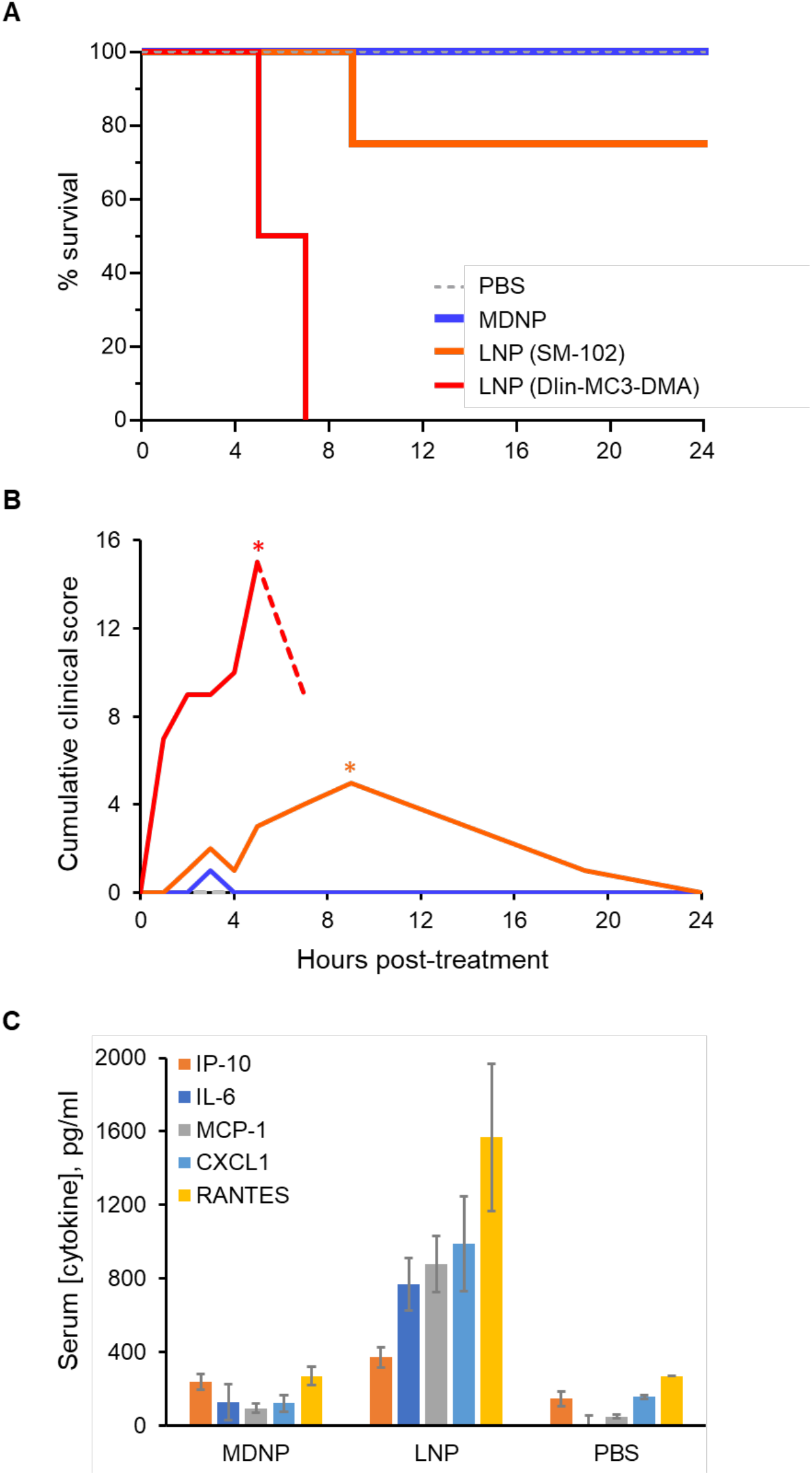
MDNP-encapsulated mRNA induces less acute inflammation than LNPs composed of DLin-MC3-DMA or SM-102. (A) BALB/cJ mice (n = 4) 6-10 weeks of age were administered 6µg of the indicated mRNA-loaded nanoparticles by IN instillation, and survival was monitored over a 24-hour period. (B) Cumulative clinical score of the same animal groups shown in (A), reflecting clinical signs of respiratory distress and abnormal behavior resulting from acute systemic inflammation according to the scoring rubric set forth in Table S1. In the DLin-MC3-DMA group, asterisk indicates time of euthanasia for 2 animals (SM-102 group) resulting in an artifactual drop in cumulative score before remaining mice were euthanized (dashed line); all 4 animals reached humane clinical endpoints by 7 hours post-treatment. Only 1 of 4 animals required euthanasia in the SM-102 group (marked by asterisk), leaving n = 3 for this group from hour 9 onward. (C) In a separate experiment modeling conventional IM vaccine administration, circulating proinflammatory cytokines were measured by ELISA in serum of IM-injected BALB/cJ mice (n = 5) 24 hours post-treatment with 5µg doses of mRNA encapsulated in MDNP or LNP composed of DLin-MC3-DMA.

**Fig S3.**
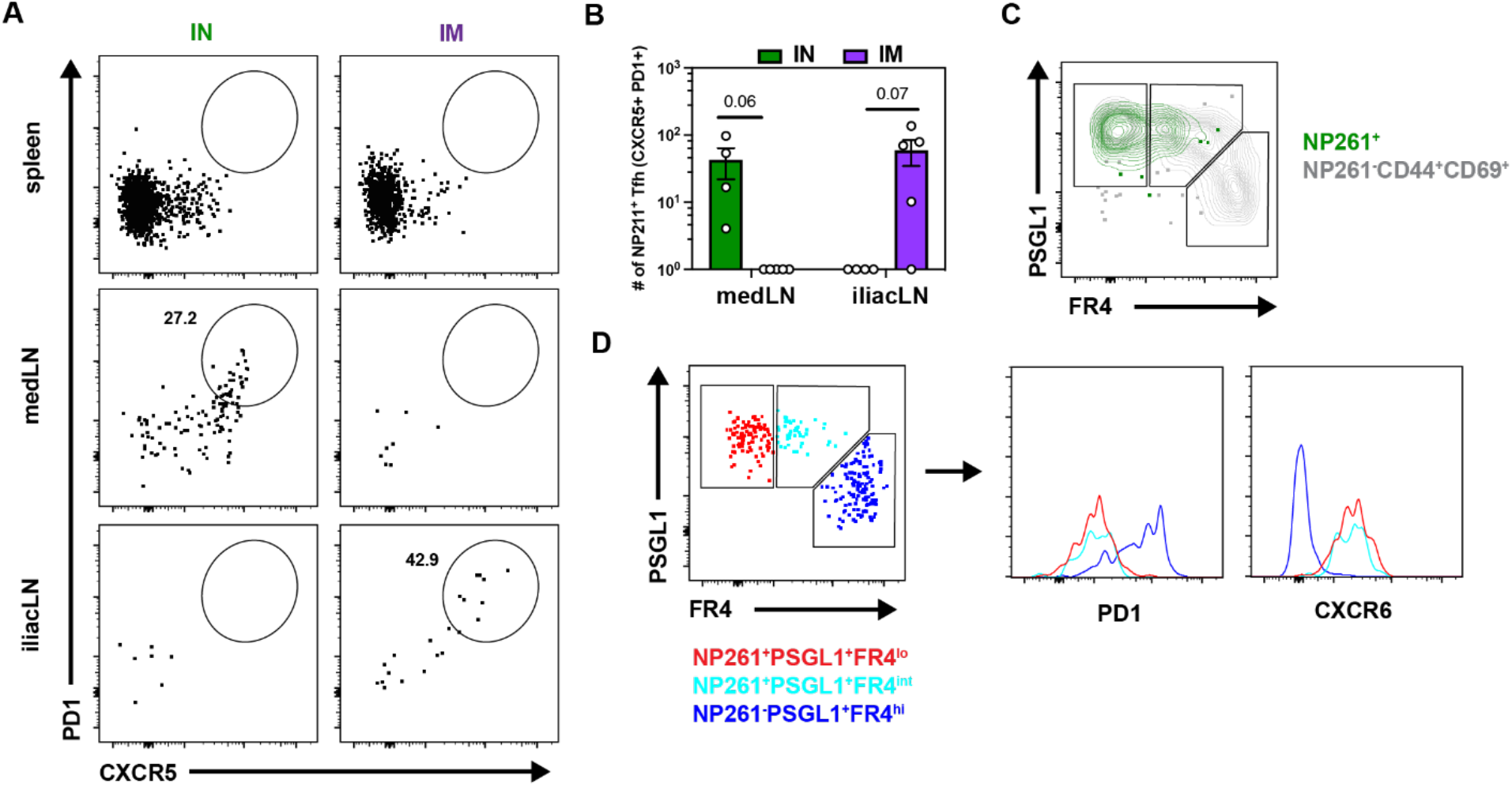
mRNA vaccination generates long-lived Tfh in the draining LN but not in the lung. (A-B) Representative flow cytometry plot (A) and quantification (B) of NP261-specific CD4+ Tfh identified by high expression of PD1 and CXCR5 in SLOs. (C) Representative flow cytometry plot of NP261+ CD4+ T cells (green) and endogenous IV-CD44+CD69+ CD4 T cell compartment (grey). (D) Representative flow cytometry plot of NP261-specific CD4+ T cell subsets in the lung tissue (left) and expression of PD1 and CXCR6 of the indicated subsets (right). Data represent N= 2 independent experiments with n = 4 to 5 mice per group. Data are shown as mean ± SEM. *P<0.05, **P<0.001, ***P<0.0001 as determined by unpaired two-tailed Student’s t test.

**Fig S4.**
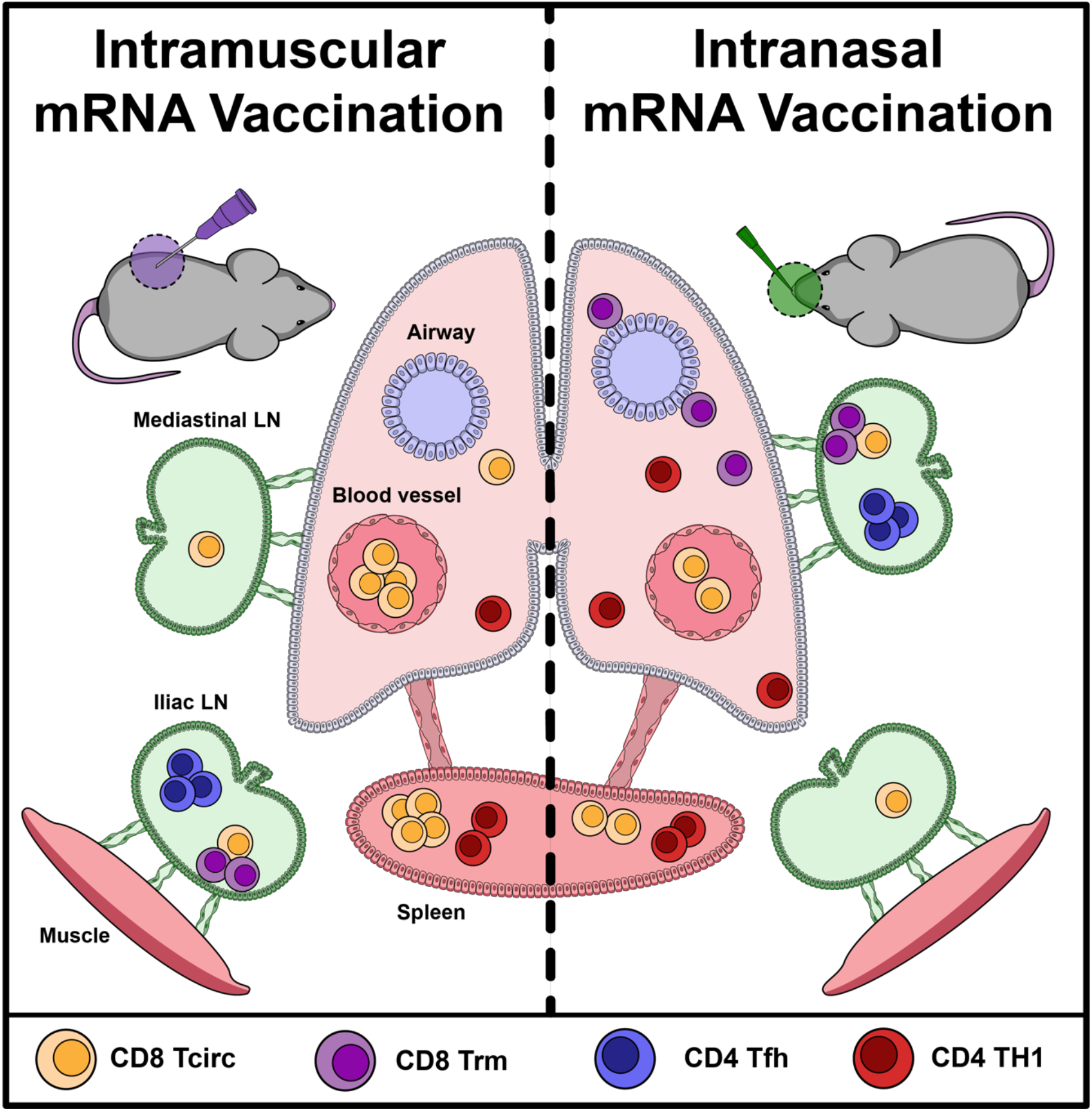
Anatomic distribution of mRNA vaccine elicited CD8 and CD4 memory T cells.

